# Subcortical recruitment dissociates isoflurane emergence from distinct wakeful states in mice

**DOI:** 10.64898/2026.05.15.725233

**Authors:** C Neiswanger, LK MacMillen, AD Murry, ER Szelenyi, J Navarrete, MHC Nota, H Lazaro, CC Shin, J Zhang, EA Apley, Y Zhang, C Diaz, KN Schneider, NL Goodwin, M Jin, SRO Nilsson, KK Ishii, GD Stuber, MR Bruchas, SA Golden, M Heshmati

**Author notes:** Centro de Investigación para Salud en América Latina, Pontificia Universidad Católica del Ecuador, Quito, Ecuador (ER Szelenyi). Co-senior authors (SA Golden, M Heshmati).

## Abstract

Emergence from general anesthesia, defined by a recovery of consciousness to the wakeful state, is a clinically consequential state transition that remains a passive process dependent on drug clearance. Despite the critical use of anesthesia, the neural circuitry underlying behavioral recovery remains poorly defined. Here, we map whole-brain neural activity during emergence from isoflurane anesthesia in mice using Fos immunolabeling, tissue clearing, and light-sheet microscopy. This approach enables unbiased quantification of neural activity at cellular resolution across the intact whole brain and supports subsequent network analysis. Rather than resembling wakefulness, emergence exhibits widespread cortical suppression alongside selective activation of discrete subcortical nuclei. This pattern of activity includes both previously implicated arousal-related regions and lesser-studied structures linked to respiratory, autonomic, interoceptive, and cerebellar function. By comparing emergence to two behaviorally distinct wakeful control states, we find that control state selection substantially shapes interpretation of whole-brain activity maps. This establishes dual-state comparisons as a broadly useful strategy for state-dependent circuit mapping. Functional network analysis further elucidates candidate central regions that strongly covary together during emergence, with the most integrated region being the ventral orbital cortex. This approach allows for targeted causal investigation, linking brain-wide circuit discovery with future hypothesis-driven mechanistic interrogation. Together, we find that emergence from isoflurane anesthesia reflects selective subcortical recruitment rather than broad global reactivation toward wakefulness.

**Significance Statement:** Millions of people undergo general anesthesia each year. While anesthetic unconsciousness is induced rapidly, emergence from altered consciousness is unpredictable. Neural mechanisms that underlie behavioral emergence remain poorly defined. Using whole-brain Fos mapping at cellular resolution, we found that emergence from isoflurane anesthesia is characterized by widespread cortical suppression alongside selective activation of discrete subcortical, autonomic, hindbrain, and cerebellar nuclei. This selective systems-level activity pattern identifies behavioral emergence as more than a simple global return toward wakefulness and highlights underappreciated neural circuitry involved in post-anesthetic recovery. Network analysis of the Fos maps further identifies candidate regions for targeted causal investigation of emergence-related regions.

## Introduction

General anesthesia induces a reversible brain state transition that is evolutionarily conserved across species, making it a powerful model for investigating the neural mechanisms underlying altered consciousness. Emergence, or recovery of consciousness and motor function after anesthesia, represents a distinct, intermediate brain state characterized by selective activation and suppression, rather than a direct return to baseline consciousness (Proekt and Kelz, 2021; Tarnal et al., 2016). There is a large literature investigating the mechanisms underlying general anesthesia-induced unconsciousness at the level of ion channels (Franks, 2008), as well as the global network level of whole-brain oscillatory activity (Brown et al., 2010). Far less is known about the circuit activation and organization during emergence at single-neuron resolution.

Studies using functional neuroimaging in humans have provided temporal definitions of brain regions activated by inhaled anesthetics (Luppi et al., 2025; Nir et al., 2022), leading to specific hypotheses that can be tested in rodents at the neural circuit level, such as the engagement of arousal circuitry by anesthesia (Brown et al., 2010; Franks, 2008). Studies in rodents have used stimulation or inhibition of arousal-related brain nuclei to uncover the functional contributions of individual brain regions to emergence (Muindi et al., 2016; Solt et al., 2014; Vazey and Aston-Jones, 2014). In these studies, brain regions are chosen *a priori* for their shared role in sleep and arousal regulation (Mashour et al., 2022; Moody et al., 2021; Vanini et al., 2020). While these insights provide significant advances in understanding emergence, they rely on assumptions that canonical arousal circuitry is the key regulator of the conscious state transition. As it would be tedious to dissect each region of the brain without an overarching framework, here we seek to provide an unbiased, holistic cellular-resolution roadmap to resolve otherwise unappreciated neural circuits related to emergence.

Here we take advantage of advances in brain-wide Fos activity mapping using light sheet microscopy to address these knowledge gaps and provide cellular resolution snapshots of neural activity within the intact mouse brain during emergence. This approach uses immediate early gene (Fos) expression as a proxy for neural metabolic activity (Morgan and Curran, 1991) and builds on existing literature using Fos-positive quantification to screen for activated brain regions in altered consciousness (Jiang et al., 2026; Madangopal et al., 2022; Renier et al., 2016; Rijsketic et al., 2023; Wasilczuk et al., 2024). By using this unbiased approach within the whole intact brain, we uncover spatially distributed, cellular-resolution neural activity patterns which could not be resolved in slice-based methods and functional neuroimaging, respectively. We use isoflurane as a pharmacologic model to examine emergence from anesthesia-induced unconsciousness due to its evolutionarily conserved effect on consciousness across phylogenetic kingdoms (Kelz and Mashour, 2019), its relatively long half-life and slow pharmacokinetic offset (Hudson et al., 2014) that encompass the timescale of Fos protein expression, and its common use in in human and veterinary medicine (Fuchs et al., 2024; Sonner et al., 2000).

To distinguish emergence-associated neural activity from general wake-related processes, we compare emergence to two behaviorally distinct wakeful control states: a home-cage condition and a context-matched open-field condition. This dual-control design enables assessment of how control state selection shapes interpretation of brain-wide activity maps while identifying convergent neural features associated with emergence across distinct wakeful baselines. We further apply network analysis to emergence-associated Fos activity patterns using a pipeline developed for brain-wide immediate early gene datasets (SMARTTR)(Jin et al., 2025) to identify coordinated activity structure and determine candidate hub regions for future causal investigation. Together, we use this approach to provide unbiased whole-brain activity mapping with circuit prioritization that defines a systems-level organization of emergence from isoflurane anesthesia.

## Results

### Behavioral modeling of the emergence state transition in mice

We examine mice in 3 conditions: (1) singly housed in their home cage (“home cage”) representative of a passive wakeful state, (2) individually exploring an open field context (“open field”) to model an active wakeful state, or (3) undergoing emergence in this open field context (“emergence”) (**Fig. 1A**, please see **Methods** for additional details). Mice in the emergence condition were induced with 2-3% isoflurane until loss of righting reflex, an established behavioral proxy for loss of consciousness in rodents (Dickinson et al., 2000). Isoflurane delivery was then discontinued to allow for the return of righting reflex (RORR), a behavioral proxy for return of consciousness. We define emergence as a post-RORR state in which subthreshold isoflurane (0.3%) was reintroduced to hold mice in a state of suspended emergence. Since Fos protein expression peaks at 90-120-min after neuronal activation, this suspended state model allowed us to capture activity during the emergence state transition (**Fig. 1B and Supplemental Movie 1)**.

**Figure 1.**
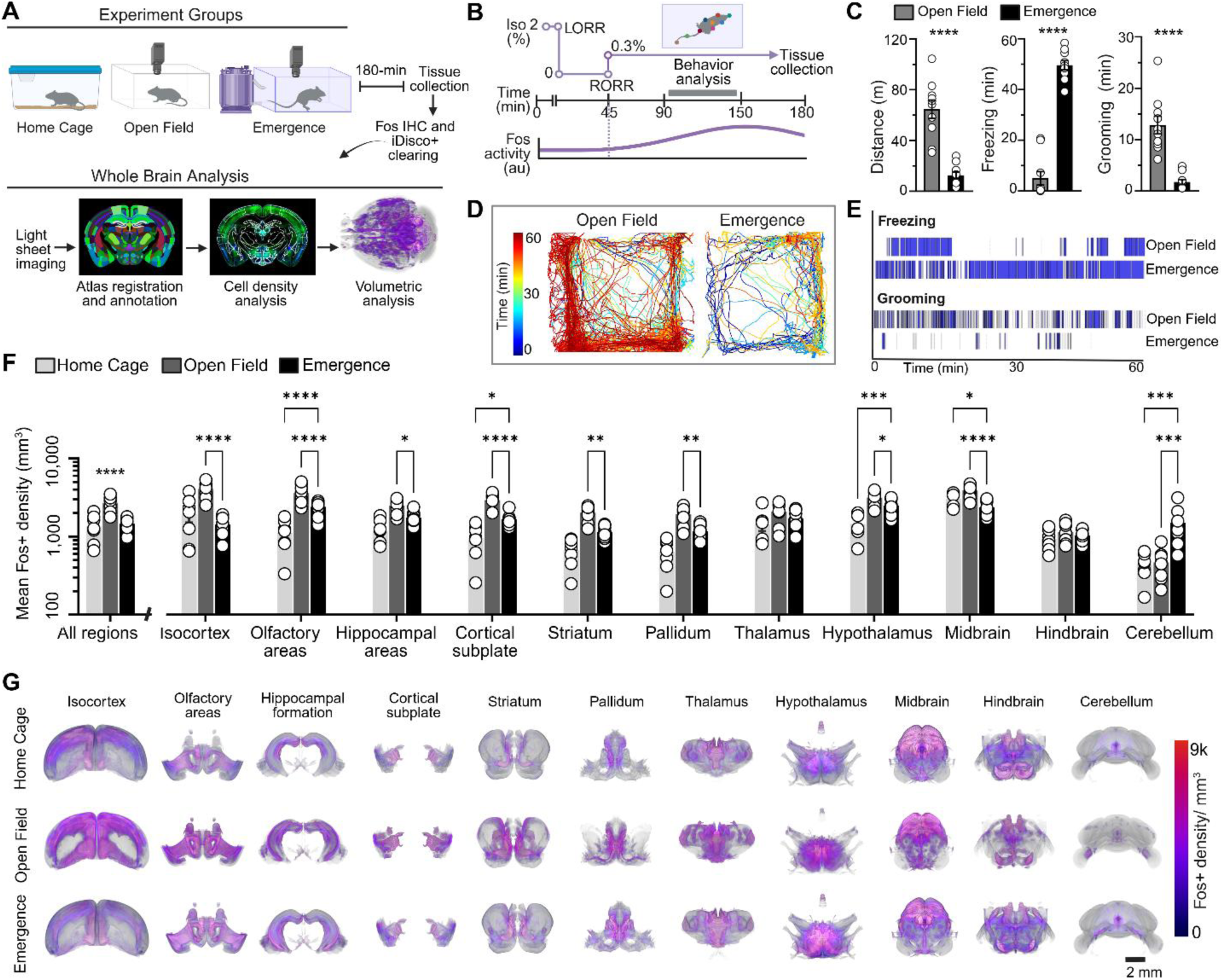
Emergence alters brainwide mean Fos density and behavior. **(A)** Experimental groups include home cage, open field, and suspended emergence. Intact brain tissue was collected after 180-min exposure to each condition, then cleared and immunolabeled for Fos using a modified iDisco+ pipeline. Below, brains were registered to the atlas and Fos expression was quantified across 914 brain subregions within 11 major divisions using cell density and volumetric analyses. **(B)** Above, timeline of suspended emergence group (LORR = loss of righting reflex, RORR = return of righting reflex); below, representation of Fos protein peak ∼90-120min after RORR. **(C)** Behavioral analysis of distance traveled (meters, m), freezing and grooming duration (minutes, min) in emergence compared to open field group (***p<0.001, t-test). **(D)** Representative distance traveled from one animal in each group with colors indicating time spent in area. A total 60-minute epoch was analyzed. **(E)** Representative gantt plots of freezing and grooming bouts from one animal in each group. **(F)** Mean Fos-positive density (-Log10 per mm^3^) in control, open field, and emergence groups across “All regions” and the 11 major brain divisions, *p<0.05, **p<0.01, ****p<0.0001, two-way ANOVA with Tukey’s multiple comparisons test. **(G)** 3-dimensional heat maps of the mean Fos+ density (0-9,000 Fos count per mm^3^) virtually sliced in the coronal plane of each major subdivision. *n* = 7 home cage, 8 open field, 9 emergence brains with the average across all samples within each group represented. Scale bar is 2mm. See **Supplemental Table 1** for detailed statistics.

We quantified behavior over a 60-min epoch using an automated supervised machine learning-based pipeline (Goodwin et al., 2024). In the emergence state, mice continued to perform behaviors such as grooming, but with significant locomotor suppression relative to the open field group (**Fig. 1B-E and Movie 1**). Distance traveled and grooming time was significantly lower in emergence, but not absent, while time spent freezing was elevated (**Fig 1C**). Representative traces from one mouse in each group of distance traveled across the 60-min epoch are shown in **Fig 1D**, along with representative Gantt plots of the freezing and grooming bouts quantified in **Fig 1E**. For details on the behavioral classifiers, please see **Methods** and **Supplemental Table 1**. We then assembled the brain wide map of regions activated during this behaviorally suspended emergence state at a fixed time point, with all brains collected 180-min following induction of anesthesia. We used iDisco+ Fos immunolabeling (Madangopal et al., 2022; Renier et al., 2014) and light sheet microscopy to image and analyze activated regions in the intact brain at cellular resolution (**Fig 1A**).

### Emergence alters brain wide mean Fos density

We analyze Fos density across 914 subregions within 11 hierarchical major brain divisions in the Unified Brain Atlas using the ClearMap open-source pipeline (Madangopal et al., 2022; Renier et al., 2016) (**Fig. 1A** and see **Methods** for further details). Results are presented as high-level overarching results within the 11 major brain divisions (**Fig. 1F-G**), followed by pairwise analyses of the 914 subregion differences between groups (**Fig. 2 and 3**). In subregion analyses, we account for multiple comparisons using the False Discovery Rate (FDR), showing significant data at an alpha threshold of 0.05 and q threshold of 0.01. We analyze significant network differences using the SMARTTR (Jin et al., 2025) pipeline, which combines whole-brain Fos quantification, atlas-based registration, and correlation-based network modeling to construct functional connectivity maps and identify putative circuitry regulating emergence in an unbiased, brain-wide manner (**Fig. 4**).

**Figure 2.**
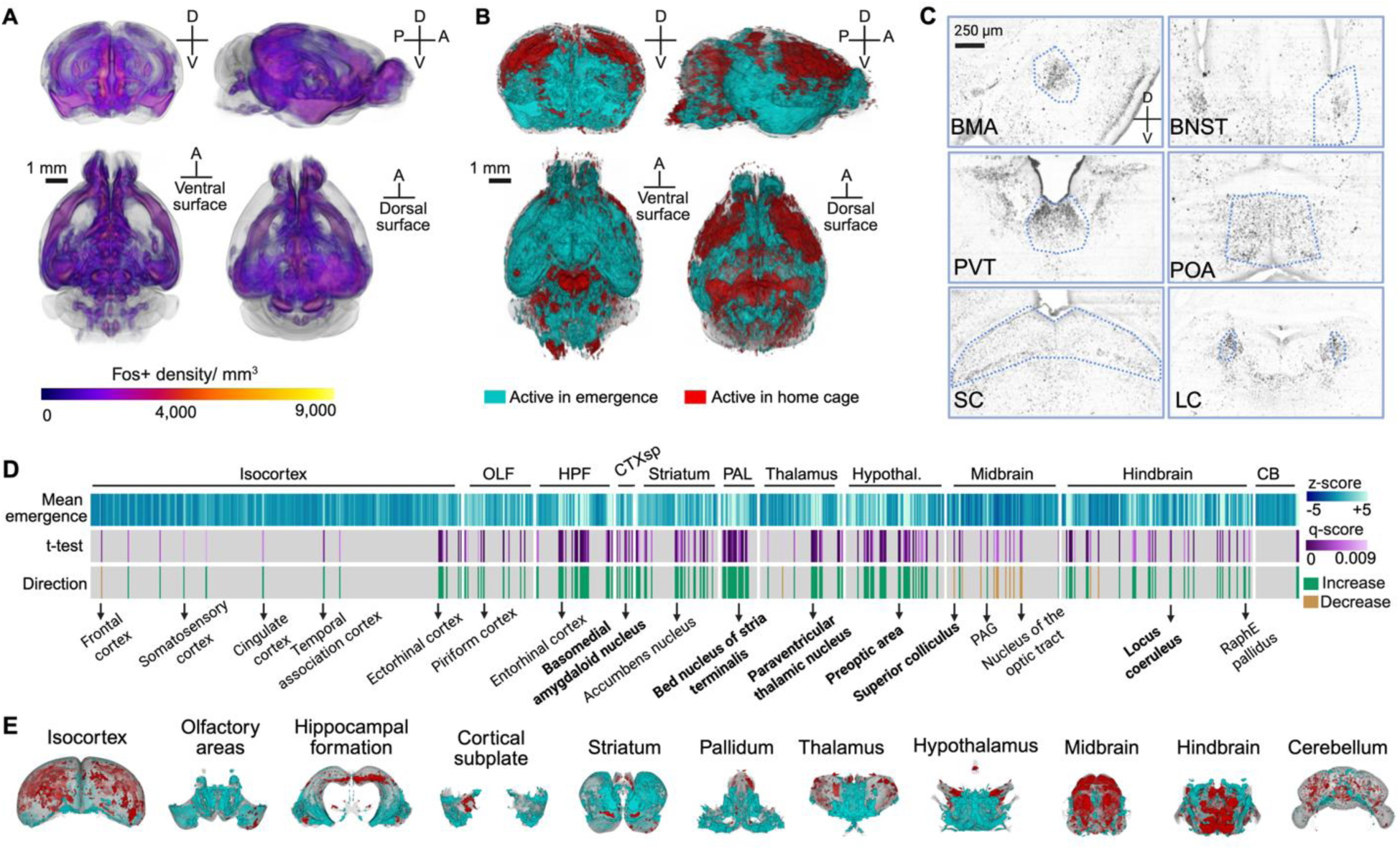
Emergence vs. home cage control pairwise comparisons localize Fos increases across spatially distributed regions. **(A)** 3-dimensional heat map rendering of mean Fos-positive density in emergence group. (**B)** 3-dimensional heat map rendering of voxelized significance in home cage control group (red) vs emergence group (cyan), scale bar 1mm. **(C)** Representative raw Fos expression in significant emergence brain regions: basomedial amygdaloid nucleus (BMA), bed nucleus of stria terminalis (BNST), paraventricular thalamic nucleus (PVT), preoptic area (POA), superior colliculus (SC), locus coeruleus (LC). **(D)** Z-score, q-score, and direction of activity for significant brain regions in emergence vs. control. t-test with false discovery rate correction of 1% (q < 0.01). Green indicates increased activity and gold indicates decreased activity. For complete list of significant brain regions See **Supplemental Table 2**. **(E)** 3-dimensional heat map representations of subregions that are significantly active in emergence (cyan) or in control (magenta).

**Figure 3.**
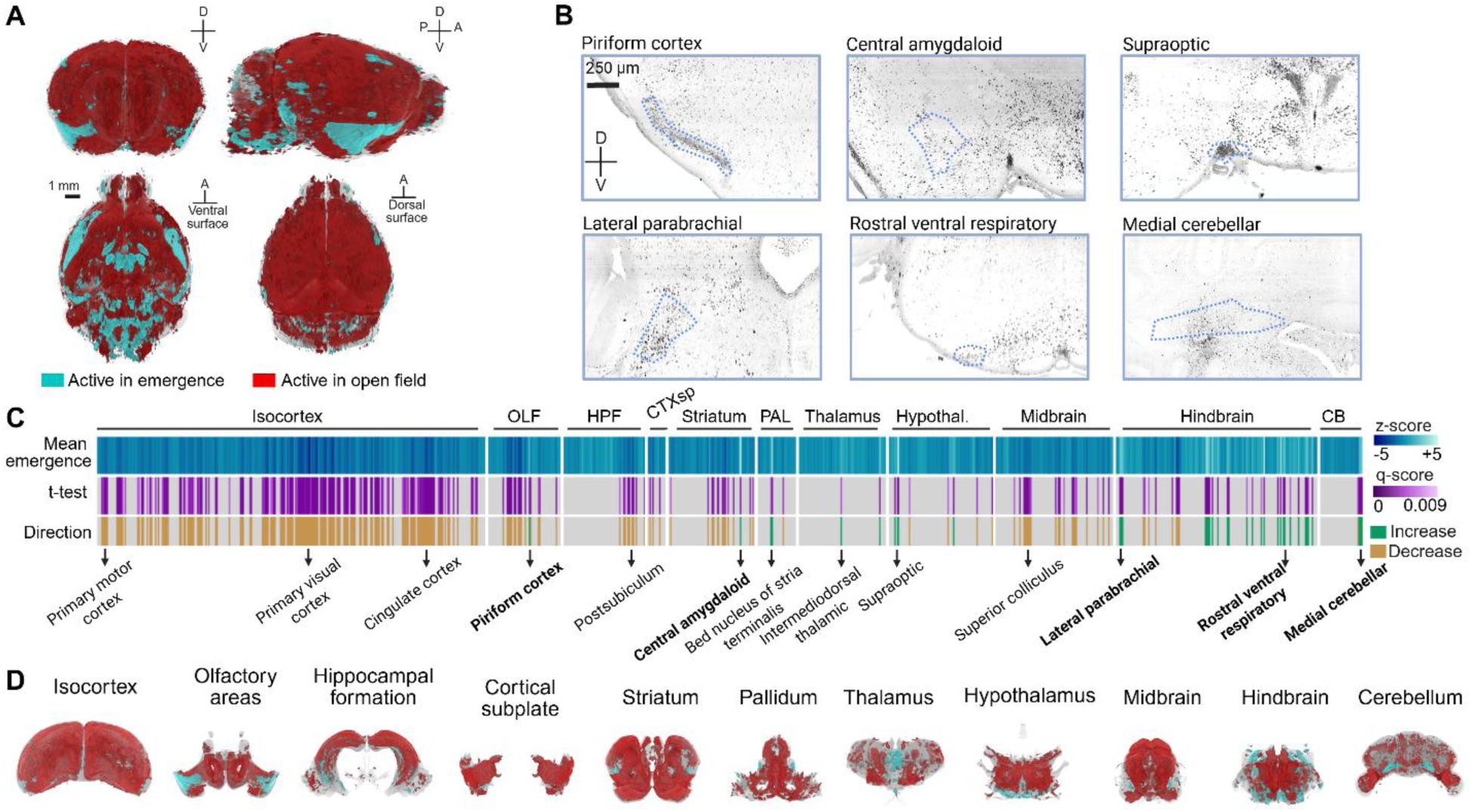
Emergence vs. open field pairwise comparisons reveal widespread cortical suppression and selective subcortical activation during suspended emergence. **(A)** Three-dimensional heat map rendering of voxelized significance in emergence (cyan) versus open field control (red) groups. Regions shown in cyan indicate significantly increased Fos expression during emergence, while red indicates increased Fos expression during open field exploration. Scale bar, 1mm. **(B)** Representative raw Fos expression in significant emergence brain regions. **(C)** z-score, q-score, and direction of activity for significant brain regions in emergence versus open field comparisons across annotated subregions. Statistical comparisons were performed using t-tests with false discovery rate correction of 1% (q < 0.01). Green indicates increased activity in emergence and gold indicates decreased activity relative to open field controls. For the complete list of significant brain regions see **Supplemental Table 3**. **(D)** Three-dimensional heatmap representations of subregions that are significantly active in emergence (cyan) or in open field controls (red) across major brain divisions.

**Figure 4.**
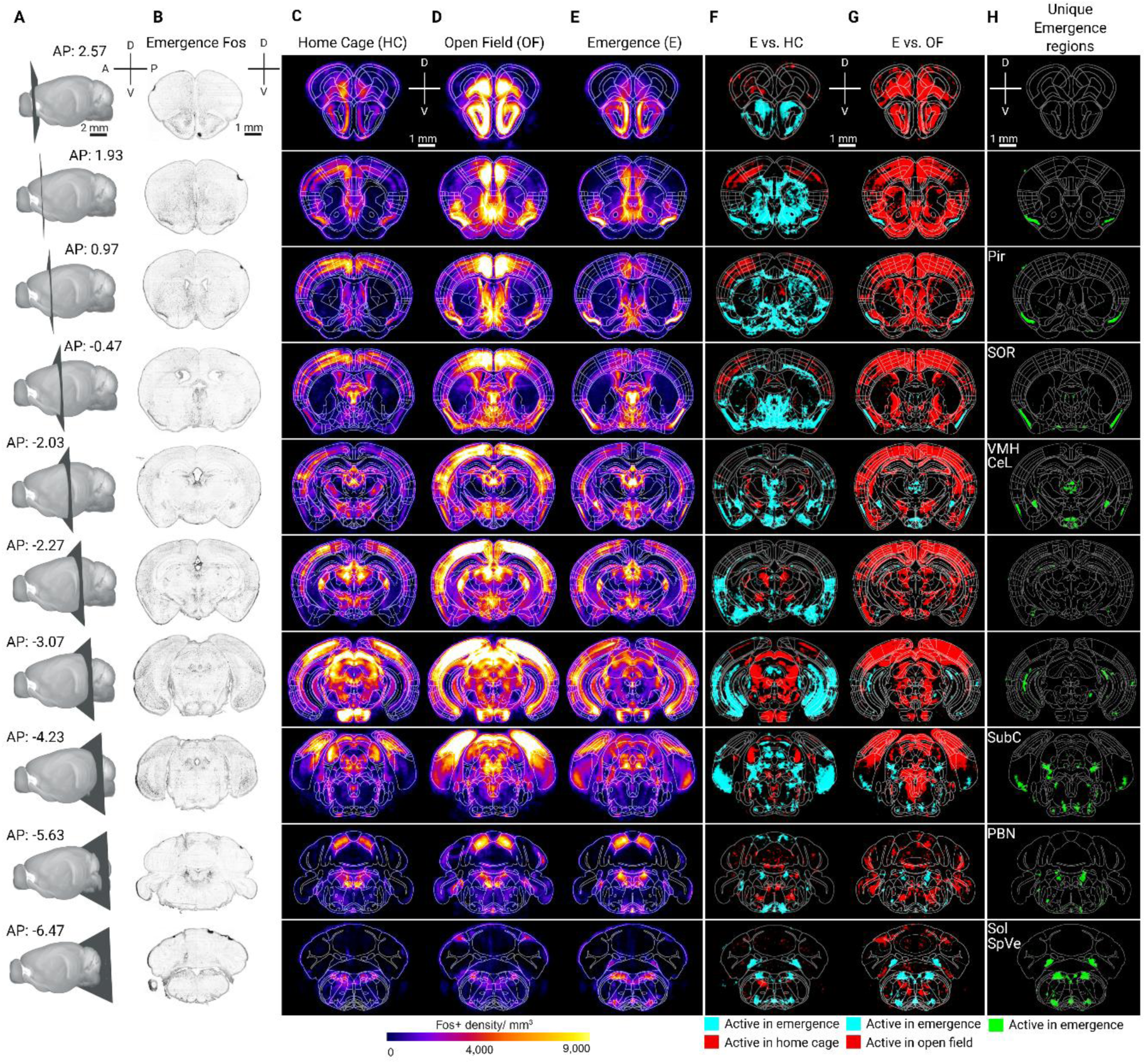
Fos activity mapping reveals brainwide signature of emergence from isoflurane anesthesia compared to home cage and open field context-matched controls. **(A)** Sagittal representation of the coronal plane shown in panel B with the anteroposterior (AP) position described in millimeters, scale bar 2mm. **(B)** Representative raw images of Fos immunolabeling in the coronal plane in one emergence sample, scale bar 1mm. **(C-E)** Coronal heat map of the mean Fos-positive density (0-9,000 per mm3) in all home cage control **(C)**, open field control **(D)**, and suspended emergence **(E)** samples at each AP position. **(F)** Coronal heat map of significant voxelized brain regions active in emergence compared to home cage controls (E vs HC) and (**G**) emergence compared to open field controls (E vs OF). Cyan blue denotes regions with significantly increased Fos in emergence, and red indicates significantly increased Fos in the control state. **(H)** Overlapping significant regions in emergence compared to both HC and OF controls. Green denotes regions significant in emergence. Scale bar is 1mm. D, dorsal; V, ventral; A, anterior; P, posterior. n = 7 home cage control, 8 open field control, and 9 emergence brains. For the complete list of significant brain regions see **Supplemental Table 2-4**

We identify distinct patterns of whole brain activity in emergence relative to the home-cage and open field control groups, as quantified by mean Fos density expression (**Fig. 1F)**. Overall mean Fos density is elevated in the open field relative to other groups across analyzed samples (1-way ANOVA, ****p<0.0001). We analyze Fos density within 11 major brain divisions: isocortex, olfactory areas, hippocampal areas, cortical subplate, striatum, pallidum, thalamus, hypothalamus, midbrain, hindbrain, cerebellum (**Fig. 1F**, 2-way ANOVA with Tukey’s test for multiple comparisons, *p<0.05). We indicate statistically significant findings in the emergence group for clarity. For full statistics and comparisons between the control groups, please see **Supplemental Table 1**.

Mean Fos density significantly decreases in the isocortex, hippocampal areas, striatum, and pallidum in emergence relative to the open field group (**Fig. 1F**, *p<0.05, **p<0.01, ***p<0.001). There are significant differences across all groups in the olfactory areas, cortical subplate, and hypothalamus, with emergence Fos density being higher than the home cage control but less than the open field control in these 3 major brain divisions (**Fig. 1F**). Relative to both control groups, mean Fos density decreases in the midbrain in emergence (**Fig. 1F**, *p<0.05,****p<0.0001). In contrast, mean Fos density increases in the cerebellum in emergence relative to both control groups (**Fig. 1F**, ***p<0.001). These quantitative differences are also observed when visually comparing anatomic hotspots of higher mean Fos density, shown as an average across all samples in each of the 3 conditions (**Fig. 1G**). Across major brain divisions, emergence was therefore characterized by a dissociated pattern of regional engagement in which broad forebrain suppression co-occurred with selective activation in specific subcortical and cerebellar systems. Together, these findings indicate that behavioral emergence is associated with selective systems-level organization rather than generalized restoration toward passive or active wakefulness.

### Emergence recruits spatially distributed subcortical ensembles relative to home cage controls

To identify region-specific activation in the suspended emergence state, we next performed pairwise comparisons of mean Fos density across all 914 brain atlas subregions. 3-D voxelized renderings of emergence mean Fos density show spatially distributed activity with subcortical hotspots (**Fig 2A**; also see **Supplemental Movie 2**). We describe statistically significant differences between emergence and the home cage control condition (**Fig 2**), followed by differences between emergence and the open field control condition (**Fig 3**).

Pairwise voxel analysis of activity in emergence vs. home cage controls reveals predominantly cortical activation in controls and subcortical activation in emergence (**Fig 2B**). Representative immunolabeling within subcortical hotspots confirms increased Fos activity in emergence (**Fig 2C**). Subregion-level analysis shows significant increases in discrete nuclei across major brain subdivisions, with areas of significantly decreased Fos predominantly in midbrain regions, such as superior colliculus (**Fig 2D; Supplemental Table 2**). Areas of increased emergence Fos activity include the basomedial amygdaloid nucleus (BMA), bed nucleus of the stria terminalis (BNST), paraventricular thalamic nucleus (PVT), preoptic area (POA), superior colliculus (SC), and locus coeruleus (LC) (**Fig 2C-D**).

3-D heatmap representations show areas of significant activation across 11 major brain subdivisions (**Fig 2E**). In isocortex, there is a significant suppression of frontal cortex in the suspended emergence state relative to home cage controls, with increased activity in all other significant cortical regions such as cingulate cortex, somatosensory cortex, and temporal association cortex (**Fig 2D-E**; see **Supplemental Table 2** for full list of significant regions).

### Emergence is characterized by suppressed cortical activity relative to open field controls

To further categorize neural activity associated with the suspended emergence state, we analyzed mean Fos density between emergence and the open field control group across the same annotated brain subregions (**Fig 3**). Open field controls were tested within the same chambers used to model the emergence state, providing a control for the environmental context (**Fig 1**). Open field exploration also represents a behaviorally active wakeful state, providing a reference for neural activity associated with ongoing locomotion and environmental engagement.

Voxelized pairwise comparisons reveal widespread regions of greater Fos expression in open field controls relative to emergence, particularly across large portions of isocortex (**Fig 3A**). Subregion analysis confirms extensive suppression of cortical Fos expression in emergence compared to open field controls, with selective activation of the piriform cortex (**Fig 3B-C**). Within subcortical areas, there is activation of central amygdaloid nucleus, lateral parabrachial nucleus, rostral ventral respiratory nucleus, and medial cerebellar nucleus in emergence (**Fig 3B-C**). Three-dimensional heat map renderings of significant voxels active in emergence relative to open field controls (cyan blue) depict the discrete olfactory, subcortical, and cerebellar areas of activation (**Fig 3D**). See **Supplemental Table 3** for a full list of significant regions.

### Subcortical regions are significantly active in emergence relative to both control conditions

Pairwise analyses identify 17 unique subregions actively recruited in emergence when compared to both control conditions (**Fig 4**). There is a significant activation of piriform cortex in emergence that is also visually prominent in raw Fos coronal images (**Fig 4B**). Remaining areas of shared significant activation are restricted to subcortical regions that include BNST and lateral parabrachial nucleus, as well as hypothalamic nuclei (see **Supplemental Table 2-4** for full list). 3-D heatmap renderings of the mean Fos density in each group further illustrate the spatial distribution of activity relative to each control group (**Fig 4C-H**). Analysis of the emergence state relative to the home cage control (**Fig 4F**) reveals widespread increases in Fos expression, including areas of frontal cortex. The emergence versus open field control comparison identifies a more restricted subset of overlapping subcortical regions at the same FDR threshold of 1% (**Fig 4G**). This pattern indicates that the open field group serves as a refinement control, narrowing the set of emergence-associated regions to those engaged outside of baseline and context-dependent wakefulness (**Fig 4H**, see **Supplemental Table 4** for a list of shared activated regions). Overall, these comparisons suggest the emergence state is dissociable from both the active wakefulness of open field exploration and the home cage wakeful state.

### Functional network analysis reveals state-dependent organization of neural activity

To understand the organization of emergence-related neural activity relative to open field controls, we analyzed each network by identifying coordinated patterns of activity across regions to detect functional network structures (Jin et al., 2025). (**Fig 5**). While the generated Fos maps show activation in individual regions, the network analysis determines how these regions covary together. Correlation matrices revealed structured patterns of co-activity across major anatomical divisions in both open field and emergence states. (**Fig 5A, B**). The open field network exhibits predominantly positive correlations across cortical and hippocampal regions, whereas the emergence network shows a broader distribution of both positive and negative correlations, including increased anti-correlation across midbrain and hindbrain regions.

**Figure 5:**
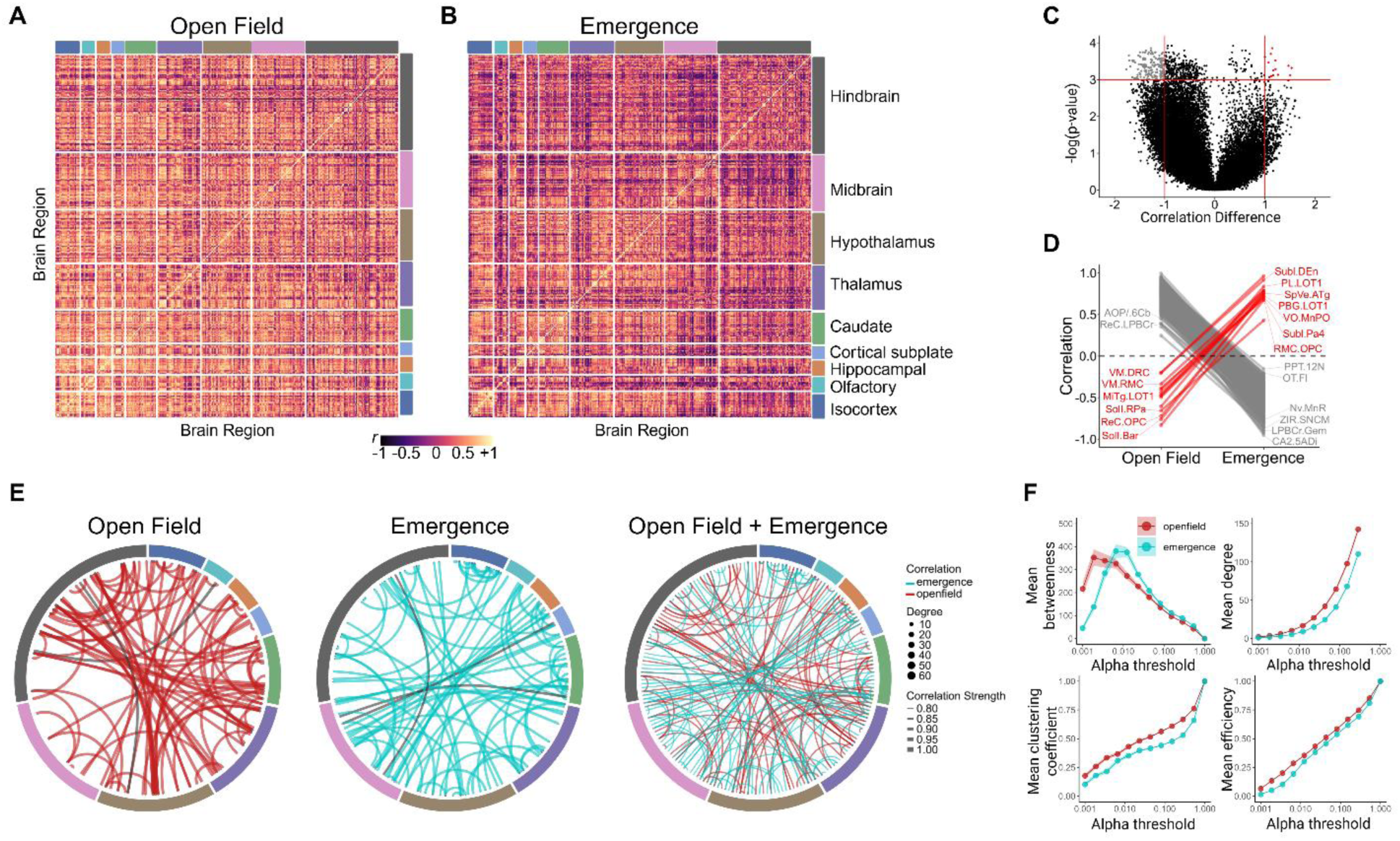
Brain-wide functional network organization differs between open field exploration and emergence from isoflurane anesthesia. Regional correlation heatmaps for **(A)** open field control and **(B)** emergence groups based on mean Fos density data for each group. Subregions are grouped by major anatomical divisions (isocortex – dark blue, olfactory – light blue, hippocampal - orange, cortical subplate - blue, caudate - green, thalamus - purple, hypothalamus - brown, midbrain - pink, hindbrain - gray). Color scale indicates correlation strength (*r,* –1 to +1). **(C)** Volcano plot showing the regional correlations that are most different between the networks. The horizontal red line denotes a significance threshold of p < 0.001. Red points in the upper right (open field < emergence) and grey points in the upper left quadrant (open field > emergence), defined by vertical red lines, indicate regional correlation differences with a magnitude shift greater than 1 between groups. **(D)** Parallel coordinate plot showing significantly positive (red) and negative (grey) regional correlations between open field and emergence conditions. **(E)** Functional networks generated by thresholding correlation matrices to retain the strongest significant positive and negative correlations in open field (left), emergence (middle), and a combined network showing the functional connectivity patterns of the individual open field control (red) and emergence (blue) functional networks. Nodes are arranged by anatomical class, and edges represent significant correlations (red/blue are positive for open field and emergence, respectively, and black are negative) **(F)** Network metrics across a range of correlation thresholds. Mean betweenness centrality (top left), mean degree (top right), mean clustering coefficient (bottom left), and mean efficiency (bottom right) for open field and emergence networks. Metrics were computed across threshold space to assess the stability of network topology. Network construction and analysis were performed using SMARTTR open-source toolkit in R (see Methods). Please see **Supplemental Table 5** for list of significant permutation results and brain region abbreviations.

We then performed a permutation analysis (see Methods) to identify functional connections (pairwise correlations) that significantly differed between states (**Fig 5C**). Plotting these significantly altered connections as a parallel coordinate plot further illustrated the directionality of change – for example, shifts from negative to positive correlations or vice versa between the open field and emergence groups (**Fig 5D).** We observe both increases and decreases in correlation strength, indicating state-dependent differences in functional connectivity.

The thresholded functional networks demonstrate distinct connectivity patterns (**Fig 5E**). The open field network exhibits dense connectivity across cortical regions and more limited connectivity in the midbrain. The emergence network shows similar overall density with more evenly distributed connectivity. A combined network visualization shows the open field and emergence networks have minimal overlap in edge structure, with largely distinct sets of connections defining each state.

We assessed global network properties across a range of correlation thresholds to evaluate possible differences in the propensity for information transfer (**Fig 5F**). Mean betweenness, reflecting the extent to which nodes act as bridges between regions, degree, representing the number of functional connections per node, clustering coefficient, reflecting the tendency of regions to form locally interconnected modules, and global efficiency, an estimate of distributed information transfer across the network, varied as a function of threshold but remained stable within each state, indicating a robust difference in network topography. Relative to open field network, the emergence network exhibited reduced clustering across thresholds, suggesting weaker local modular organization and less efficient integration of tightly interconnected regional subnetworks. This reduced local coordination is consistent with prior models describing emergence as a more stochastic and unstable transitional brain state.

### Community analysis identifies state-dependent network hubs that connect cortical and subcortical nodes

We next asked if there was a coordinated organization amongst the connections most significantly changed between the open field and emergence groups. To assess this, we performed community detection analysis, which revealed 8 distinct communities composed of anatomically distributed regions, spanning both cortical and subcortical areas (**Fig 6**). These communities are essentially larger units of functional changes between groups that allow for visualization of central nodes. Several nodes exhibit a high degree, including the supraoptic nucleus (SOR), subcoeruleus nucleus (SubCA), ventral orbital cortex (VO), and the solitary nucleus (SolC). The SOR, SubCA, and SolC are central nodes within communities enriched for connections stronger in open field compared to emergence, whereas the VO was the prominent hub within a community characterized by connections increased in emergence relative to the open field state. Importantly, hub status in the communities reflect coordinated covariance structure rather than absolute Fos density, indicating that VO may function as an integrative node during emergence despite relatively modest mean activation levels.

**Figure 6.**
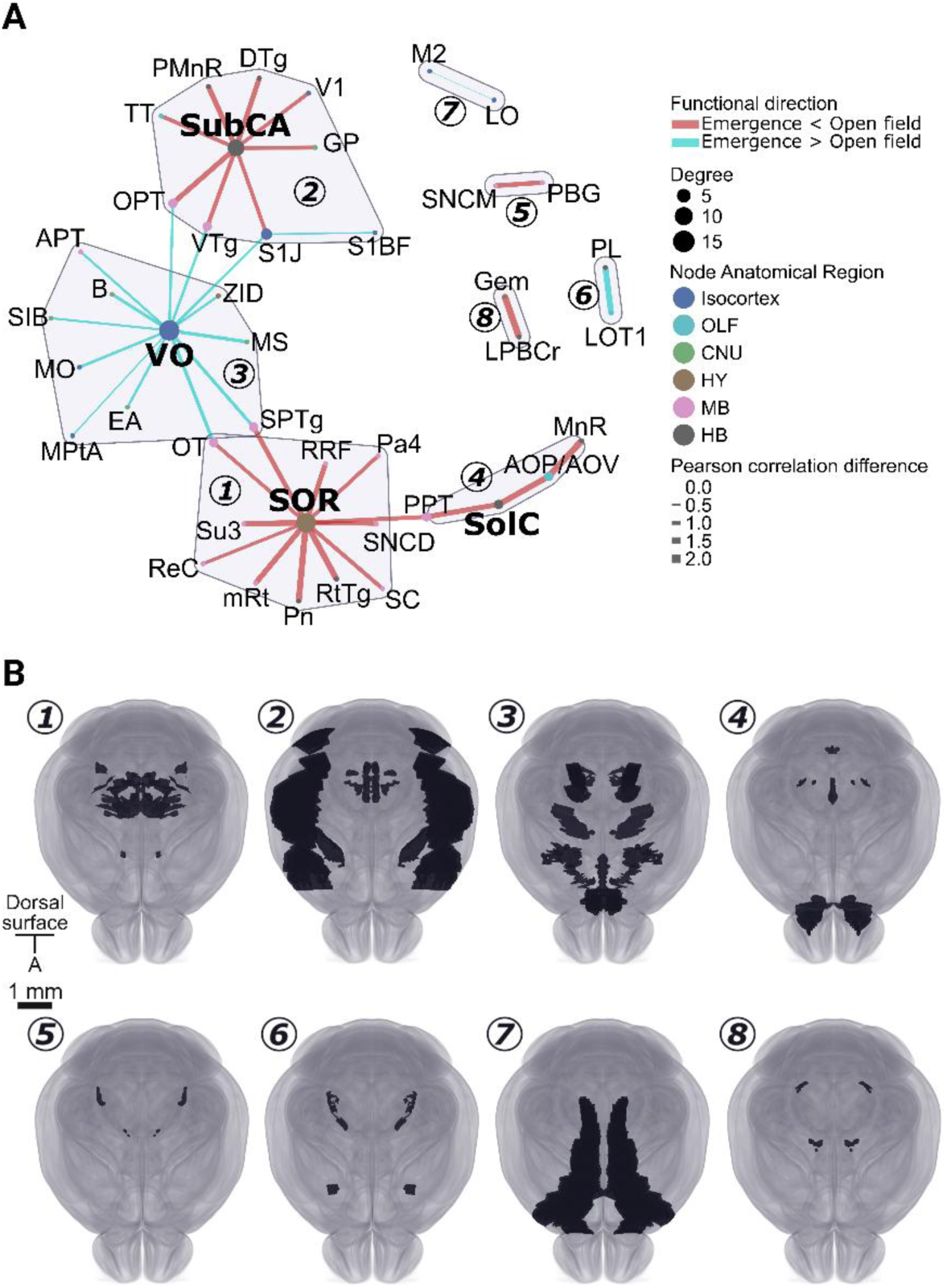
Community structure of functional network differences in emergence vs. open field control state. **(A)** Functional difference network generated by plotting significant permuted regional correlation differences as edges (lines) and the regions they connect as nodes (circles). Edge color denotes directionality of the difference (emergence < open field, red; emergence > open field, blue), and node size reflects degree. Edge shading represents the magnitude of Pearson correlation differences. Convex hull polygons delineate unique communities. Communities are labeled and key nodes indicated as follows: 1: SOR (supraoptic nucleus, retrochiasmatic part); 2: SubCA (subcoeruleus nucleus, alpha part); 3: VO (ventral orbital cortex); 4: SolC (solitary nucleus, commissural part); 5: SNCM–PBG (substantia nigra compact part, medial tier; parabigeminal nucleus); 6: PL–LOT1 (paralemniscal nucleus; nucleus of the lateral olfactory tract 1); 7: M2–LO (secondary motor cortex; lateral orbital cortex); 8: Gem–LPBCr (gemini hypothalamic nucleus; lateral parabrachial nucleus, crescent part). Node colors indicate anatomical classification. **(B)** Anatomical distribution of regions within each community (1–8) mapped onto a dorsal brain template. Scale bar, 1 mm. See **Supplemental Table 6** for full list of abbreviations and significant community nodes.

Four smaller, less connected communities associated with emergence and open field, such as secondary motor cortex (M2) and lateral orbital cortex (LO), are segregated, while the four larger communities are interconnected through shared nodes and cross-community edges. This organization reflects distinct but interacting network modules, with state-dependent differences in connectivity patterns distributed across subcortical and cortical regions.

Several nodes identified within these communities, including hypothalamic, parabrachial, and pontine regions, overlap with brain systems previously implicated in arousal regulation, sleep-wake transitions, and anesthetic state control. In contrast, the emergence-associated prominence of cortical regions such as the ventral orbital cortex suggests coordinated recruitment of higher-order integrative circuitry not typically emphasized in traditional arousal-centered models of emergence. Together, these findings indicate that recovery from isoflurane anesthesia involves distributed state-dependent network reorganization spanning autonomic, limbic, and cortical systems rather than isolated activation of canonical wake-promoting nuclei.

## Discussion

Mechanisms by which the brain restores consciousness following anesthesia remain incompletely understood. In addition, no active strategies exist to accelerate recovery of consciousness in routine clinical practice (Vincent et al., 2025). As a result, emergence from anesthesia remains an unpredictable transition to consciousness, with complications such as respiratory and cardiac events, delayed cognitive recovery, uncontrolled pain, agitation, and delirium (Cascella et al., 2020; Eckenhoff et al., 1961). There is an urgent need to better define neural mechanisms engaged by emergence that may advance the development of therapeutic strategies to facilitate improved recovery.

Here, we present a brain-wide, cellular resolution resource for identifying state-dependent neural activity in emergence from isoflurane anesthesia. By applying intact brain clearing with Fos immunolabeling and light-sheet microscopy, this approach enables unbiased quantification of activity across the intact brain, overcoming limitations of regionally targeted or slice-based analyses. Importantly, this framework allows for the identification of spatially distributed neural interactions at cellular resolution within the intact brain that would not be captured using traditional approaches.

Comparative analyses across two distinct control states reveal that emergence from anesthesia represents a distinct brain state defined by selective subcortical recruitment and cortical suppression. Directionality of activity across subregions embodies a distributed pattern of network-level modulation. While several subcortical regions exhibit robust increases in activity, other regions show significant decreases, suggesting emergence is characterized by coordinated reorganization rather than global reactivation. Notably, the interpretation of emergence-related activity patterns is influenced by the choice of control state. Comparisons to home cage controls emphasize recruitment of subcortical structures relative to baseline, whereas comparisons to open field controls indicate widespread cortical suppression relative to active wakefulness. These findings highlight that control state selection is a critical factor shaping conclusions from brain-wide immediate early gene activity patterns.

Across analyses, we identify significant activation of the olfactory areas, hindbrain and cerebellum in emergence. Existing literature supports modulation of arousal state by olfactory stimuli, with specific studies examining noradrenergic and slow-wave sleep regulation (Geng et al., 2025; Jia et al., 2016; Manabe et al., 2011; Yamaguchi, 2017). Piriform cortex encodes higher order features of odors in both mice and humans (Kehl et al., 2024). While studies have examined olfactory detection and amnesia under anesthesia, further research is needed to evaluate the relationship between piriform cortex modulation and conscious state transitions.

Network reorganization in the emergence state highlights distinct configurations of circuit activation and suppression, with predominant subcortical activation. The supraoptic nucleus (SOR) and the subcoeruleus nucleus (SubCA), responsible for neuroendocrine regulation and REM sleep regulation, respectively, emerged as main hubs of network connectivity in the direction of the open field control state, with relative suppression in the emergence state. These areas play key roles in sedation and consciousness, including modulating recovery of consciousness (Zhang et al., 2025). The SOR is implicated in neuroendocrine regulation of arousal and contains a population of known anesthesia-activated cells (Jiang-Xie et al., 2019).

Ventrolateral and median preoptic areas of the hypothalamus also control anesthetic sedation and emergence in a cell-type and region-specific manner (Moore et al., 2012). The subCA is relatively understudied in emergence, though known to play a role in sedation and REM sleep modulation as a component of the pontine reticular formation (Fraigne et al., 2015; Schott et al., 2023). A neighboring region, the locus coeruleus (LC), is a known population of noradrenergic cells that regulate wakefulness and emergence from anesthesia (Du et al., 2025; Vazey and Aston-Jones, 2014). LC is identified as significantly activated when compared to home cage control, though it does not emerge as significantly correlated within the emergence network, suggesting it is an independent contributor to emergence. Causal mechanistic studies targeting LC, SOR and SubCA are needed to disentangle the contributions of these specific subregions from neighboring areas in emergence.

In addition, the ventral orbital cortex (VO) displayed high-degree connectivity in the network characterized by widespread, though relatively low-magnitude, increases during emergence compared to the open field state. The VO plays a role in reward value representation and integration of sensory with internal state information to shape decision making (Kringelbach, 2005). High centrality of VO in the emergence network and its strong correlation with other cortical subregions indicates a critical role of recruiting decision-making circuitry during recovery of consciousness. This activity pattern supports a framework in which higher-order processing is recruited early during emergence, as was shown in a recent study in humans (Mashour et al., 2021). This further suggests recovery of consciousness may be defined within global neuronal workspace theory, supporting previous studies using anesthesia to manipulate consciousness (Mashour et al., 2022, 2021, 2020). The interaction of these cortical circuits with subcortical limbic systems processing emotional reward valence may influence states of hyperarousal or agitation that are observed in patients during emergence from anesthesia (Heshmati and Bruchas, 2022).

Emergence- and open field-biased subnetworks form partially segregated communities that are connected by a limited number of low-degree nodes, indicating weak integration between state-specific network configurations. This organization suggests that emergence does not recapitulate the network structure observed during natural behavior but reflects a transitional state in which distinct functional configurations coexist with constrained interactions. Such configurations may reflect incomplete reintegration of distributed networks or persistence of state-dependent circuitry during recovery from anesthesia, supporting models of hysteresis and neural inertia in emergence (Proekt and Kelz, 2021).

A few limitations should be considered when interpreting these findings. Fos-based mapping reflects activity-dependent transcription at a snapshot in time rather than direct neuronal firing and does not distinguish between excitation of principal neurons and recruitment of inhibitory cell populations. As a result, Fos-positive activation in this dataset may reflect an activation of GABAergic inhibitory populations that result in net suppression of brain areas.

There is also limited temporal resolution. We adapted a behavioral model to suspend emergence across the Fos protein expression window (Herrera and Robertson, 1996) and collected a snapshot in time. We also did not include a neurophysiologic or electroencephalogram (EEG) measure of the brain state in emergence. Prior work demonstrates substantial variability in EEG dynamics and state-switching during emergence state transitions (Akeju et al., 2014; Purdon et al., 2013; Stone et al., 2025; Warnaby et al., 2017). As an alternative, we apply a behavioral definition of emergence in this dataset. In addition, our statistical approach in pairwise comparisons was intentionally conservative (FDR correction with q < 0.01) and may exclude potentially relevant regions as false negatives while more confidently identifies the most strongly activated regions. Despite the limitations, these data provide a robust, unbiased framework for identifying candidate neural circuits engaged by emergence from isoflurane anesthesia. This supports future hypothesis-driven causal interrogations of the identified brain regions.

## Materials and Methods

### Mice

C57BL/6J male and female mice, aged 2-3 months, were used for all experiments. All animals were housed in 12h/12h reverse light cycle with free access to standard food and water and tested midday during the dark/awake phase. All experiments were approved by the National Institute of Health and the University of Washington Institutional Animal Care and Use Committee (IACUC).

### Experiment conditions

2-3-month-old C57BL/6J male and female mice are used in all groups (See **Fig 1A**, **Experimental timeline)**. *n* = 10 open field, 10 emergence (5 female). Due to tissue damage that occurred during brain processing, 8 open field (3 female) and 9 emergence (4 female) intact brains are included in Fos analysis.

(1) *Open field* control mice were habituated to a 30×30cm box daily for 3 days. On the test day, they were placed in the box with video recording throughout the 180-min trial. They were perfused at the end of the session.
(2) *Emergence* mice were habituated to the same 30×30cm box daily for 3 days. The test box is adapted with inlets to administer and scavenge isoflurane anesthesia, along with gas analyzer sampling of the isoflurane concentration. Test boxes were serially connected to one isoflurane vaporizer, and isoflurane concentration within the chambers was recorded using a gas analyzer with continuous sampling throughout the experiment. On the test day, mice were individually induced with 2-3% isoflurane and placed on the back in the 30×30cm test box to confirm loss of righting reflex. Isoflurane concentration was gradually reduced to 0% delivery until return of righting reflex occurred, approximately 45-min later. Mice were then maintained at 0.3% isoflurane, as recorded by the gas analyzer, and allowed to move freely within the 30×30cm chamber with concurrent videotaping of activity for an additional 150-min prior to transcardial perfusion. The total duration of all group trials was 180-min followed by immediate perfusion.
(3) The *Home cage* control group is a re-analysis of a previous dataset (Lazaro et al., 2025). *n* = 7 (3 female). *Home cage* control mice were singly housed and habituated to the procedure room daily under red light conditions/awake phase for 3 days. The total duration of single housing was less than 1 week. Mice remained singly housed in their home cages for the duration of the 180-min test period and did not undergo additional handling or anesthetic exposure. They then underwent transcardial perfusion for tissue collection under isoflurane anesthesia promptly at the end of the 180-min period.

### Perfusion and post-fixation

All animals were deeply anesthetized with isoflurane and transcardially perfused with phosphate-buffered saline (PBS) and 10% formalin. The brains were dissected and post-fixed in formalin for 24 hours at 4°C and stored in PBS containing 0.02% sodium azide prior to immunolabeling and clearing. Brains were collected for clearing and imaging.

### iDisco+ clearing and Immunolabeling

A modified iDisco+ protocol was used to first immunolabel and then clear intact brain samples (see Madangopal, Szelenyi et al 2022; Renier et al 2016). Briefly, samples underwent dehydration using an increasing gradient series of 20%, 40%, 60%, 80%, 100% methanol (MeOH) and deionized water (45 minutes each wash), followed by delipidation with 66% dichloromethane and 33% MeOH (8 hours plus overnight). Samples were washed in 100% MeOH and bleached in 5% hydrogen peroxide in MeOH overnight. Samples were next rehydrated using a decreasing gradient series of MeOH: 80%, 60%, 40%, 20%, 0%, 45 min each wash at room temperature, followed by washing in PBS with 0.2% Triton-X 100 (PBS-T) Samples were next incubated in permeabilization buffer containing PBS-T, 1.4% glycine, and 20% dimethyl sulfoxide (DMSO) (2 days at 37°C), and then placed in blocking buffer containing PBS-T, 6% normal donkey serum (NDS) and 10% DMSO (3 days at 37°C). They were washed in PBS with 0.2% Tween-20 and 10ug/ml heparin (PTwH buffer), followed by incubation in the primary antibody - 1:2500 rat monoclonal anti-Fos IgG (Synaptic Systems 226 017) in PTwH with 3% NDS (7 days). Samples were washed in PTwH and incubated in the secondary antibody – 1:500 AlexaFluor 647 Fab2 Donkey anti-rat IgG in PTwH with 3% NDS (2 days). After immunolabeling, samples were dehydrated in an increasing gradient series of 20-100% MeOH in water followed by delipidation in 66% dichloromethane and 33% MeOH and washed in 100% dichloromethane to remove remaining MeOH. Samples were last placed in 100% dibenzyl ether for clearing and refraction index matching overnight.

### Light sheet microscopy

Cleared intact mouse brains were imaged using uniform axial resolution light sheet microscopy (SmartSPIM, LifeCanvas Technologies) with a 1.6X resolution objective (LifeCanvas LCT 1.6.02). Select high resolution images were taken with a 3.6X objective (Thor Labs TL4x-SAP). Samples were mounted in horizontal orientation using a custom sample holder submerged in dibenzyl ether. Autofluorescence excitation at 488nm was acquired in conjunction with Fos immunolabeling at 647nm in separate 2×2 tile scans.

### ClearMap analysis

We processed whole brain light sheet-acquired images using a modified version of the ClearMap open-source toolkit. We first digitally re-sliced images in the coronal plane and registered each sample to the Unified Brain Atlas. We then calculated Fos-positive cell counts using automated cell detection within two levels of hierarchy defined by the Unified Brain Atlas; (1) the 11 major anatomical divisions of the brain and (2) 914 minor subregions within those higher-level anatomical divisions that each no longer split into further distinct subregions as defined by the atlas. To ensure accuracy within the hierarchical relationships, we manually verified that there were no overlapping spatial footprints, and no double counting of any regions, following each subregion branch until the end. For the 11 major brain divisions’ mean Fos density analysis, we used a two-way ANOVA with the within-subject factor of region and the between-subject factor of treatment group (Home Cage, Open Field, Emergence) followed by Tukey’s post hoc multiple comparisons at alpha <0.05. **See Supplemental Table 1 for detailed statistics**. For subregion pairwise analysis, we used pairwise t-tests with FDR correction for multiple comparisons at q <0.01. We normalized the raw Fos cell counts for each region by calculating z-scores relative to the mean Fos count of the control group used (Home Cage or Open Field) to account for regional volume differences. Z scores for each region of interest were calculated using the formula z= (x-μ)/σ, where x represents the raw Fos count of the treatment group, μ represents the mean Fos count of the control group, and σ represents the SD of the control group for each region. z-scores are converted into -Log10 values for visualization purposes. We provide comprehensive raw count data, z-scored data, and statistical results for both high-level and stop-level analyses in Supplementary Tables. Analysis was performed using custom python code and GraphPad Prism software.

### Whole brain visualization and voxel-cluster analysis

3-dimensional whole brain image stacks were visualized using Arivis (Zeiss) scientific image analysis platform. Figures created using Arivis (Zeiss) and BioRender.

### Network and Community analysis

Functional network analysis was conducted using the open-source SMARTTR R script package (Jin et al., 2025). Whole brain Fos positive cell densities as quantified across 914 anatomically defined regions by ClearMap were normalized and used to compute pairwise Pearson correlations across animals within each condition, generating region-by-region correlation matrices. Statistical significance of correlations was assessed using asymptotic p-values from one-sample t-tests and corrected for multiple comparisons using FDR (q < 0.05). Condition-dependent functional connectivity differences between emergence and open field networks were identified by calculating for each regional pair and evaluating using a permutation-based approach. Null distributions were generated by randomly shuffling group labels across animals (50,000 iterations) and recomputing correlation differences at each iteration. Empirical p-values were derived from these distributions, and significant edges were defined at p < 0.001.

Differential networks were visualized with brain regions as nodes and edge weights corresponding to the magnitude of correlation differences. To ensure robustness, networks were evaluated across a range of statistical thresholds. Graph theoretical properties were computed at the nodal level and summarized to obtain global network metrics. All analysis code is available at https://mjin1812.github.io/SMARTTR/. See **Supplemental Tables 5-6** for detailed statistics.

### Animal tracking and behavioral analyses

Behavioral analyses were performed using methods available in the SimBA API (Goodwin et al., 2024). For details, see https://simba-uw-tf-dev.readthedocs.io/. Body parts pose-estimated in SLEAP were smoothed (Savitzky-Golay, 300ms) and missing data was interpolated (nearest reliable detection). After computing pixel-to-metric conversion factors, distance travelled and travel paths were computed using the pose-estimated center key point proxying location.

Grooming and freezing were scored as previously described (Lazaro et al., 2025). Briefly, for grooming we used a random forest model with custom features dependent primarily on the distribution of head area, body length, and snout movement within sliding temporal windows of 0.25–2s. For freezing, we used a heuristic model where freezing was scored as present in frames where velocity (computed from the mean movement of the nape, snout, and tail-base key points) fell below 5mm/s over the preceding 3s. We performed t-tests using GraphPad Prism. See **Supplemental Table 1** for detailed statistics.

## Author Contributions

MH and SAG conceptualized and directed all aspects of the study. CN and MH wrote the first draft of the manuscript. ERS, MJ, KKI, GSS and SAG developed whole brain clearing, imaging and analysis pipelines. ADM, JN, YZ and MH handled mice for behavioral studies. ADM and MH collected whole brain samples. ADM, ERS and JN cleared, immunolabeled and imaged whole brain samples. CN, LKM, ADM, ERS, MJ, SAG and MH analyzed whole brain datasets. HL, CCS, JZ, EAA, NLG, and SRON analyzed behavioral video datasets. CN, MHCN, CD, KNS, KKI, GSS, MRB, SAG and MH revised the manuscript. All authors approved the final manuscript.

## Acknowledgements

Funded by the National Institute of General Medical Sciences R35GM146751 (to M.H.), Foundation for Anesthesia Education and Research (to M.H.), National Institute on Drug Abuse UG3DA053802 (to S.A.G.) and R37DA033396 (to M.R.B.). We also thank funding from the National Institute of General Medical Sciences F32GM163495 (to C.N.), Washington Research Foundation Fellowship (to E.R.S.), National Institute of Mental Health F31MH133295 (to J.N.), National Institute of Health F31MH125587 (to N.L.G.), and the University of Washington Center for Excellence in Neurobiology of Addiction Pain and Emotion P30DA048736. Figures created using Arivis (Zeiss) and BioRender software.

## Supplemental Tables

Table 1: Individual Fos density values for major anatomical regions

Table 2: Emergence vs home cage control pairwise comparisons of 914 subregions

Table 3: Emergence vs open field control pairwise comparison of 914 subregions

Table 4: Overlapping subregions with significantly increased Fos expression in emergence

Table 5: Network analysis permutations in emergence vs open field control

Table 6: List of community analysis nodes and edge values in emergence vs open field control

## Movies

Movie 1: Behavioral classification of grooming (left) and freezing (right) showing the same mouse in the suspended emergence state.

Movie 2: Brainwide mean Fos density heat map of emergence group virtually re-sliced in the coronal plane.

